# Predicting CTCF-mediated chromatin loops using CTCF-MP

**DOI:** 10.1101/259416

**Authors:** Ruochi Zhang, Yuchuan Wang, Yang Yang, Yang Zhang, Jian Ma

## Abstract

The three dimensional organization of chromosomes within the cell nucleus is highly regulated. It is known that CTCF is an important architectural protein to mediate long-range chromatin loops. Recent studies have shown that the majority of CTCF binding motif pairs at chromatin loop anchor regions are in convergent orientation. However, it remains unknown whether the genomic context at the sequence level can determine if a convergent CTCF motif pair is able to form chromatin loop. In this paper, we directly ask whether and what sequence-based features (other than the motif itself) may be important to establish CTCF-mediated chromatin loops. We found that motif conservation measured by “branch-of-origin” that accounts for motif turn-over in evolution is an important feature. We developed a new machine learning algorithm called CTCF-MP based on word2vec to demonstrate that sequence-based features alone have the capability to predict if a pair of convergent CTCF motifs would form a loop. Together with functional genomic signals from CTCF ChIP-seq and DNase-seq, CTCF-MP is able to make highly accurate predictions on whether a convergent CTCF motif pair would form a loop in a single cell type and also across different cell types. Our work represents an important step further to understand the sequence determinants that may guide the formation of complex chromatin architectures.

## Introduction

Three dimensional organization of the chromosomes in the human genome is critically important for understanding the principles of gene regulation and disease mechanisms [1–4]. Recent high-throughput mapping methods such as Hi-C [5, 6] and ChIA-PET [7, 8] have revealed that higher order genome organizations harbor more complex global chromatin interactions than we previously thought. One of the most intriguing examples involves the architectural protein CCCTC-binding factor (CTCF). In addition to serving as insulator, CTCF is known to have the capability of forming chromatin loops especially with the cohesin protein complex [3, 6, 9]. Through CTCF depletion, a recent work directly demonstrated that CTCF has critical roles in forming chromatin loops and establishing insulation between topologically associating domains (TADs) [10]. Importantly, global mapping data of chromatin interactions based on higher coverage Hi-C and ChIA-PET both found that the majority of CTCF binding sites at chromatin loop anchor regions are in convergent orientation [6, 8], suggesting that motif orientation may also play key roles in establishing CTCF-mediated loops. Indeed, a recent study using CRISPR-cas9 showed that the change of orientation of specific CTCF binding site could have major impact on long-range chromatin architecture and gene regulation [11].

However, several important questions related to CTCF chromatin loops remain elusive. For example, not all the CTCF binding sites in the genome form chromatin loops. Even if the CTCF protein binds to a CTCF binding site on the genome in a particular cell type, it may not be able to form chromatin loops with other sites. Furthermore, for a pair of convergent CTCF motifs that are both bound by the CTCF protein in a particular cell type, it also does not always form chromatin loop. Therefore, the following question remains: are there other features in addition to CTCF motifs, especially sequence-based ones, in the genomic context that may be important to establish CTCF-mediated chromatin loops? Recent observations showed that various epigenetic marks may provide clues to predict CTCF loops [12]. However, the roles of sequence-based features are still unclear. In this paper, we directly tackle this question. We are primarily interested in revealing the contribution of sequence-based features that are predictive for the formation of CTCF-mediated chromatin loops without leveraging much help from functional genomic signals. The motivation is to decode potential instructions already encoded in our genome that govern chromatin organization. Such knowledge is particularly important when we interpret mutations in human disease genomes. Specifically, we address the following questions:

a. What are the main sequence level differences between CTCF motifs that form loops and those that do not form loops?
b. For a certain cell type, can we train a model to predict whether a pair of convergent CTCF motifs bound by CTCF would form a chromatin loop in that cell type?
c. Can we train a model based on existing cell type(s) to predict whether a pair of convergent CTCF motifs bound by CTCF would form a chromatin loop in a new cell type?

We developed a series of computational methods to approach these questions. In particular, we designed a new machine learning algorithm based on word embedding [13] to address Questions (b) and (c) (see Methods). Our main contribution is three-fold: (1) We identified important sequence-level features that can help distinguish CTCF motifs that form loops and those that do not. We found that motif conservation measured by “branch-of-origin” [14] that accounts for motif turn-over in evolution is a very informative feature. (2) We developed a new machine learning algorithm, called CTCF-MP, based on word2vec and boosted trees to demonstrate that sequence-based features have the capability to predict if a pair of convergent CTCF motifs would form a loop. (3) We further demonstrated that we can build an effective model based on data from existing cell types to predict chromatin loops formed by convergent CTCF motif pairs in a new cell type. We believe our work represents an important advancement in understanding the principles of CTCF-mediated chromatin loops with the potential to decode information embedded in the genomic sequences that guide the formation of complex chromatin architectures. The source code of CTCF-MP can be accessed at: https://github.com/ma-compbio/CTCF-MP.

## Results

We first compared chromatin interaction data from Hi-C and CTCF ChIA-PET in GM12878 and decided to focus on the ChIA-PET data to analyze CTCF-mediated chromatin loops in this work. In GM12878, we identified 92,808 CTCF loops from ChIA-PET data generated in [8]. For all 112,430 CTCF motifs that we identified in the human genome (see Methods), 32,312 (28.7%) of them completely overlap with the loop regions defined by ChIA-PET, where 85.1% of these motifs are in CTCF ChIP-Seq peak regions (+/- 50bp of the ChIP-seq peak summit). In addition, for a pair of loop regions where each has a unique CTCF motif, 67% of these paired CTCF motifs are in convergent orientation. These results are consistent with the previous observations from Hi-C and ChIA-PET [6, 8]. In the Hi-C data from GM12878, we identified 12,559 CTCF motifs in chromatin interaction loops and 74.7% of the motifs overlap with CTCF ChIP-Seq peaks. For all the CTCF motifs in the genome that overlap with CTCF ChIP-Seq peaks in GM12878 (38,590), 71.3% of them are involved in ChIA-PET defined loops but only 24.3% of them are within Hi-C defined loops from [6]. As discussed in [8], the specific enrichment in CTCF ChIA-PET experiments is likely to be the reason that led to more detailed map of CTCF-mediated loops. Here we therefore focus on the ChIA-PET data to analyze CTCF-mediated chromatin loops. We call the CTCF motifs that are within chromatin loop regions “loop motifs”; otherwise, we call them “non-loop motifs”.

### More ancient CTCF motifs are more likely to be involved in chromatin loops

We first explored the association between the evolutionary conservation of CTCF binding motifs and their involvement in CTCF loops. We started by using the mammalian phyloP scores [15] to look at sequence conservation at base-pair level in CTCF motifs bound by CTCF (based on ChIP-seq data) in four different cell types (GM12878, K562, HeLa, and MCF7). We found that CTCF motifs in chromatin loops overall have much higher phyloP scores than non-loop motifs (Fig. S1A). The average phyloP score of loops is generally three times higher than non-loops. In particular, the more conserved positions in the CTCF core motif PWM tend to be much more conserved in loop motifs than non-loop motifs. Specifically, the average phyloP score on position 4,5,7,10,13, and 14 for loop motifs is above 0.8 (Fig. S1A); these positions have been shown to have important roles in zinc finger binding [16].

The base-pair level conservation analysis led us to further explore the connection between CTCF motif conservation and its loop-forming capability. Transcription factor (TF) binding site turn-over events are prevalent in cis-regulatory sequence evolution [17, 18]. We previously developed a model to quantify the TF binding site conservation by taking turn-over into account [14], with which we assign the emergence of a lineage-specific TF binding site in the human genome to a particular branch in the mammalian phylogeny (i.e., “branch-of-origin”). We calculated the branch-of-origin of CTCF loop motifs and non-loop motifs using the approach in [14] (see Methods). We found that there is a significant difference in branch-of-origin between the two types of CTCF motifs (Fig. S1B). More ancient CTCF motifs are more likely to form loops. On all the ancestral branches older than the primate common ancestor (Fig. S1B), there were more loop motifs emerging as compared to non-loop motifs. These results suggest that the CTCF motifs involved in CTCF-mediated chromatin loops are more conserved evolutionarily than the motifs that do not form loops. We also showed that the branch-of-origin score is more informative as compared to phyloP (see Supplementary Results) and such information provides extra predictive power in addition to the CTCF ChIP-seq signals (see later section and Table 2).

### Overview of CTCF-MP – A new algorithm for predicting CTCF loops

Next, we developed a machine learning approach to predict whether a pair of convergent CTCF motifs can form a chromatin loop. Fig. 1 illustrates the workflow of our algorithm, named CTCF-MP, which can be summarized into four steps. (1) We generated positive and negative samples based on CTCF ChIA-PET and CTCF ChIP-seq data from a given cell type. It is important to note that in CTCF-MP we focused on convergent CTCF motif pairs that are bound by CTCF (i.e., within CTCF ChIP-seq peak). As discussed earlier, the majority of CTCF loop motif pairs show convergent orientation. If we consider all CTCF motif pairs patterns in the same dataset without removing non-convergent motif pairs, the performance of the prediction could be strongly biased because it would be easy for the classifier to distinguish positive samples from negative ones by simply using the motif pair directionality as the most important feature. (2) We developed a word2vec model (see Methods) using CTCF binding motif and its surrounding genomic sequence as input. Word2vec is a popular word embedding model in natural language processing. It reduces the dimensionality of words but keeps useful information of relationship between words. Here, we utilized this model to encode DNA sequences into continuous vectors as one of the features, which had better performance than traditional *k*-mer frequency features (see Table 2 later for details). (3) Features for the boosted trees classifier consider various sources, including the word2vec model we trained, additional features (including branch-of-origin, distance between the motif pair, motif occurrence frequency in the window region, and GC content), as well as CTCF ChIP-seq and DNase-seq signals. (4) We trained a classifier based on boosted trees to evaluate our predictions on whether a pair of convergent CTCF motifs form a loop, for both same cell type prediction and cross cell type predictions. The algorithmic details of CTCF-MP are discussed in the Methods section.

**Figure 1:**
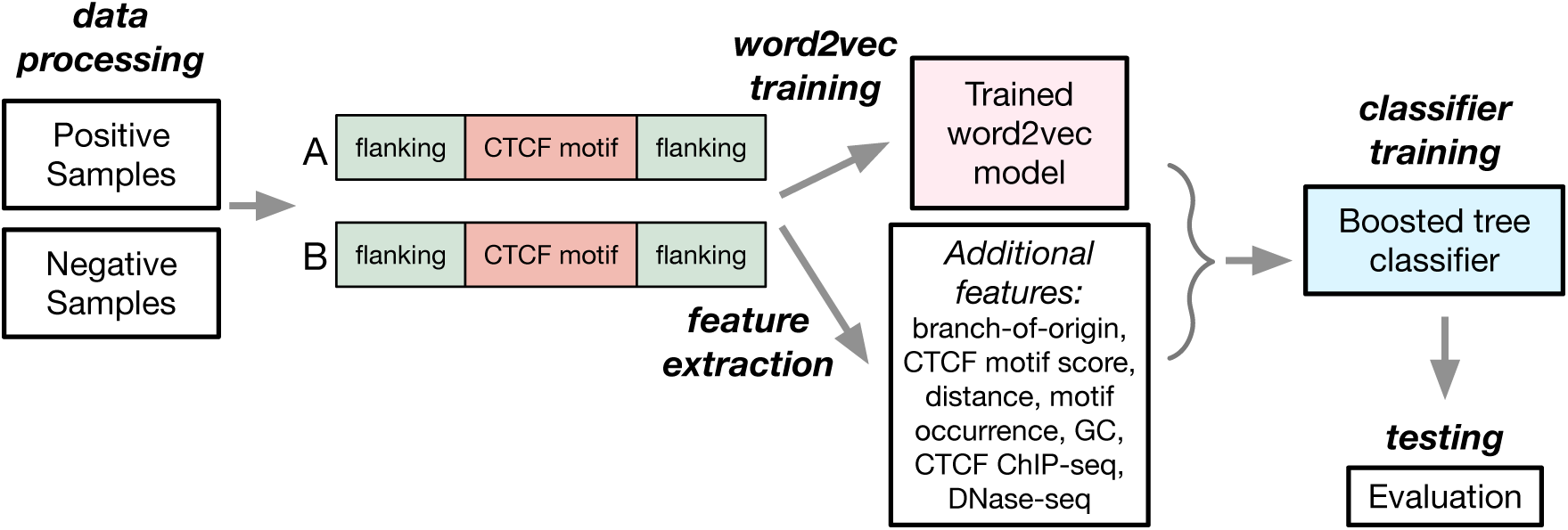
Overview of the CTCF-MP algorithm.

### CTCF-MP can predict loops formed by convergent CTCF motif pairs with high accuracy

We evaluated the trained classifier based on boosted trees in CTCF-MP to distinguish interacting CTCF motif pairs with non-interacting ones. While numerous machine learning techniques are designed for the classification problem, we chose some of the widely used algorithms and made a comparison first. Methods were evaluated through 10-fold cross-validation and measured by multiple metrics. All of the tests were conducted with balanced data from GM12878 (both positive and negative sample sizes are 21,301) and the results are in Fig. S2. Here we balanced the dataset by sampling negative data to be the same amount of positive ones with matching distance. To be more specific, for each positive sample, we selected a non-duplicated negative sample with similar distance between two CTCF motifs. We found that boosted trees had the best performance and was therefore chosen as the classifier for further analysis. Boosted trees achieved 95.5% AUROC and 95.1% AUPR, suggesting overall strong performance of the boosted trees classifier used in CTCF-MP. An example is shown in Fig. 2 to illustrate the contributions of some features used in CTCF-MP and the prediction performance. As shown in the figure, most of the CTCF loops (track “Loops in GM12878”) are between CTCF pairs that are more conserved (more ancient than the primate common ancestor). Also, CTCF-MP accurately predicts most of the CTCF loops with fewer false positives and false negatives.

**Figure 2:**
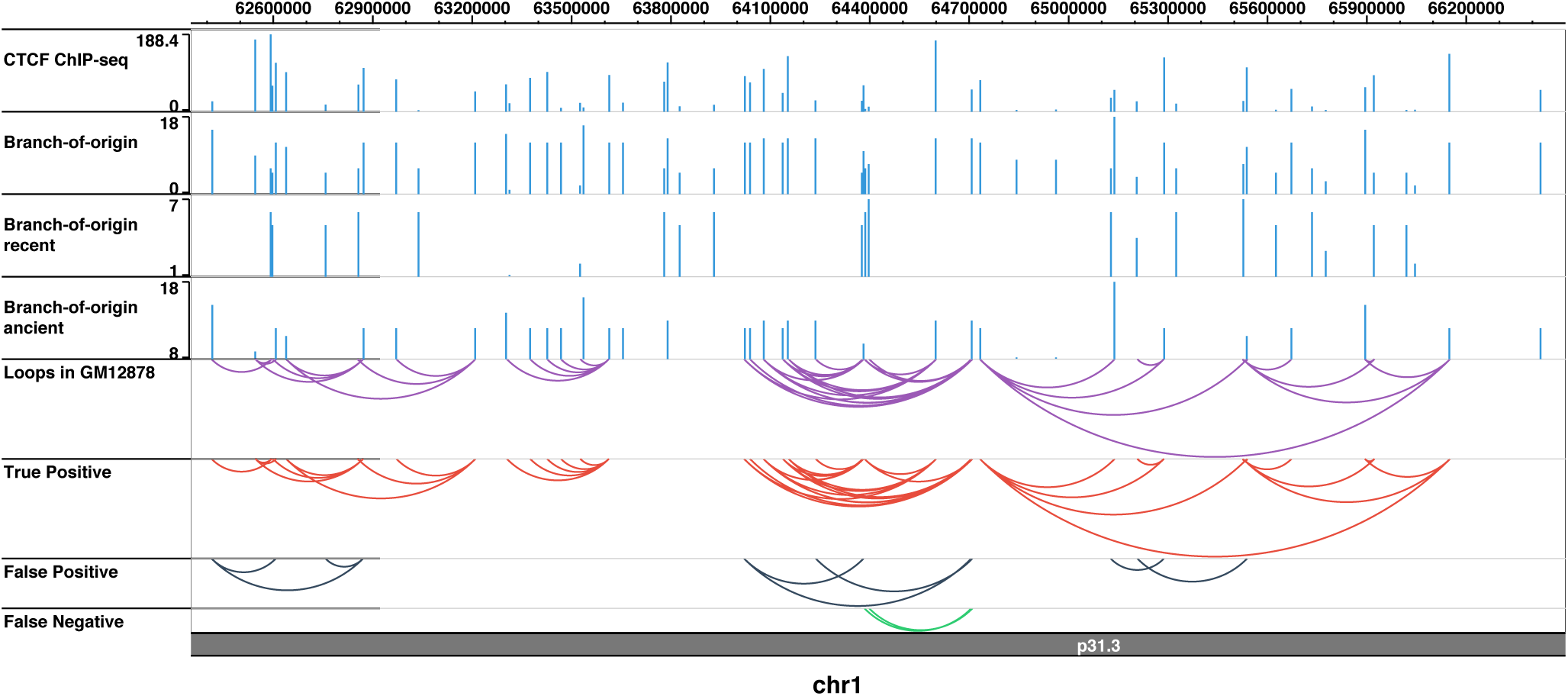
An example of the prediction from CTCF-MP. In this region (chr1:62.4Mb-66.4Mb), we visualize some features used in CTCF-MP as individual tracks, chromatin loops in GM12878 from ChIA-PET, and the predictions from CTCF-MP. “CTCF ChIP-seq” track shows the ChIP-seq peak value for the CTCF motifs in GM12878. “Branch-of-origin recent” refers to the CTCF motifs derived after the primate common ancestor and “Branch-of-origin ancient” refers to the ones older than the primate common ancestor. “Loops in GM12878” are the CTCF loops based on ChIA-PET in GM12878. “True Positive/False Positive/False Negative” are the predictions made by CTCF-MP.

To test whether the classifier is robust for different cell types, we repeated the evaluation in other three cell types: K562, HeLa, and MCF7 (both positive and negative sample size: 7,969, 9,506, 13,240, respectively, for these three cell types). The dataset and features of these cell types were generated and extracted following the same procedure for GM12878. We again used 10-fold cross validation for evaluation (see Table 1). We found that overall CTCF-MP can predict CTCF loops well across cell types, where it has the best performance on GM12878, and even in the worst case it achieves 90.3% AUROC.

**Table 1:**
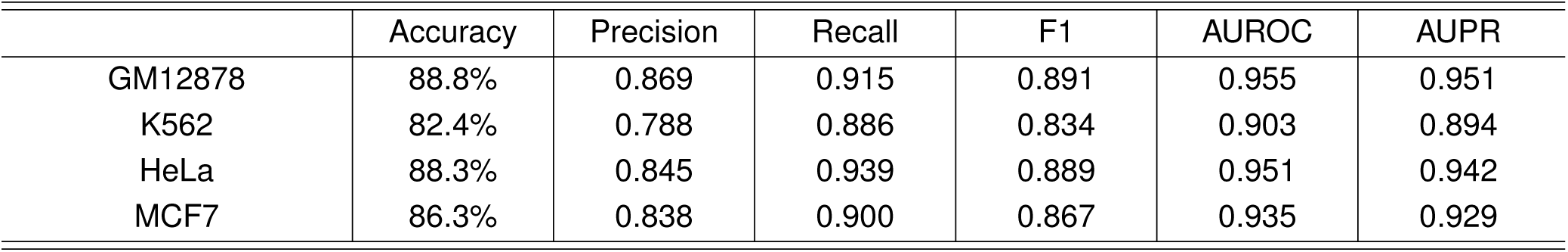
Evaluation results when applying CTCF-MP to predict loops from convergent CTCF motif pairs.

|  | Accuracy | Precision | Recall | F1 | AUROC | AUPR |
| --- | --- | --- | --- | --- | --- | --- |
| GM12878 | 88.8% | 0.869 | 0.915 | 0.891 | 0.955 | 0.951 |
| K562 | 82.4% | 0.788 | 0.886 | 0.834 | 0.903 | 0.894 |
| HeLa | 88.3% | 0.845 | 0.939 | 0.889 | 0.951 | 0.942 |
| MCF7 | 86.3% | 0.838 | 0.900 | 0.867 | 0.935 | 0.929 |

We then asked whether the performance of CTCF-MP varied between facultative (i.e., more cell type specific) loops and more constitutive loops. We grouped the convergent CTCF loops in GM12878 by the number of occurrences of each loop in all four cell types used in this study. We then calculated the prediction accuracy from cross-validation for each group. Note that since we only calculated the accuracy of positive samples, it is equivalent to the recall score in the cross-validation test. The results are summarized in Fig. 3. We found that, as expected, CTCF-MP performs better for the constitutive loops as compared to more facultative ones. For those CTCF loops that appear in all four cell types, the accuracy for those reaches 97.6%. However, even for the ones that only show up in one cell type, CTCF-MP can still achieve very high accuracy (>87%).

**Figure 3:**
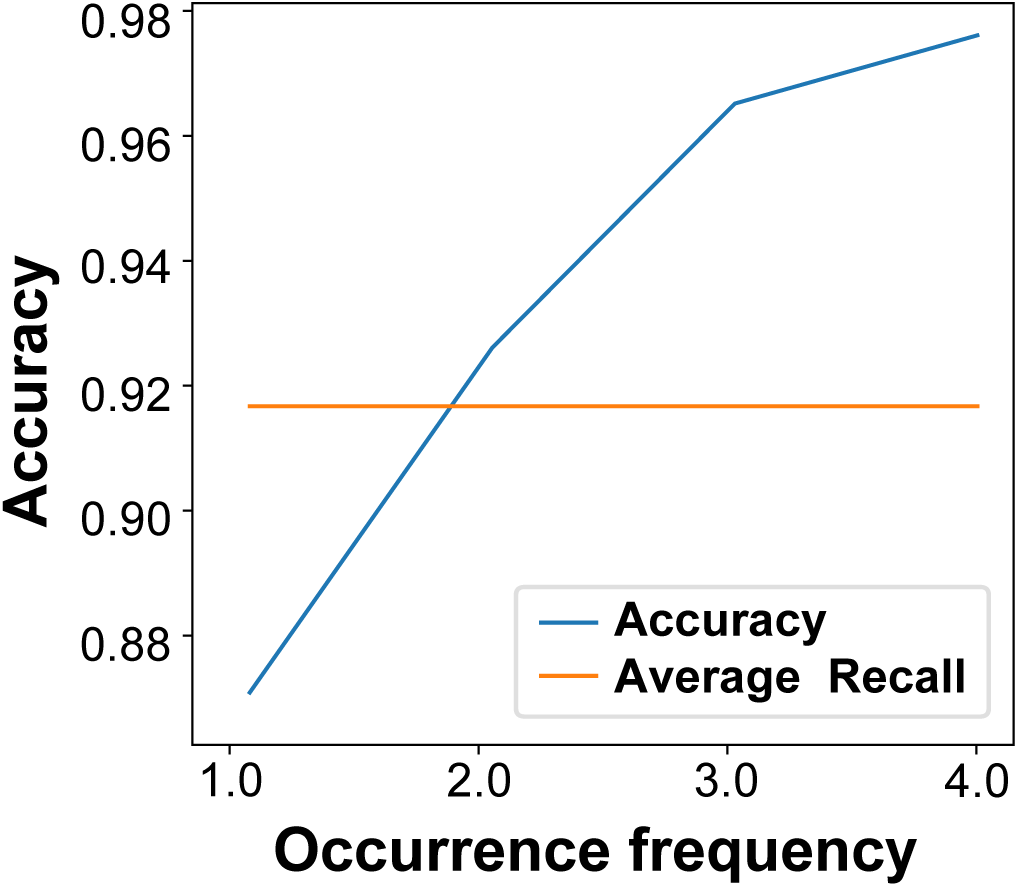
Performance of CTCF-MP over constitutive vs. more facultative loops. “Occurrence frequency” indicates the times a loop appears in four cell types.

We did additional evaluation to see if CTCF-MP’s performance would change if the distance between paired CTCF motifs changes. We grouped the datasets (both positive and negative samples) by their distance based on their genomic coordinates and calculated the average accuracy for these groups. As shown in Fig. S3, we found that the performance does not have strong correlation with distance, showing that CTCF-MP has overall strong power regardless of distance.

Overall, these results demonstrated that CTCF-MP can train an effective model to accurately predict loops formed by convergent CTCF motif pairs in a single cell type. We also found that it has strong performance even in highly imbalanced data (Supplementary Results).

### CTCF-MP can predict loops formed by convergent CTCF motif pairs in a new cell type

We then asked if we can train a CTCF-MP classifier based on existing cell type(s) and predict CTCF loops in a new cell type. We generated dataset following the same procedure described above, trained the model with dataset from one cell type, and used dataset from another cell type as testing data. Here we require that positive samples in both training and testing datasets are cell type specific loops (between the two cell types). The negative samples are also not shared between training and testing to make sure that training and testing are completely separate. However, we remark that it is possible that the same pair of CTCF motifs is positive in one cell type but negative in the other, which increases the difficulty of this cross cell type prediction task.

We found that CTCF-MP trained with data from one cell type can accurately predict CTCF loops that are specific in another cell type (see Fig. 4). In each off-diagonal entry in the figure, the number shows the AUROC for using data from one cell type (cell1) to train and then test on data from another cell type (cell2). In the entries on the diagonal, the AUROC is from cross-validation when training and testing were performed on the same cell type. As mentioned above, CTCF-MP tends to perform better on more constitutive loops as expected. However, it is interesting to observe that it also achieved high AUROC for predicting cell type specific loops. In Fig. 5 we show one example of the cross cell type predictions. It shows that even cell type specific loops can be accurately predicted by CTCF-MP. This suggests that the CTCF pairs that form chromatin loops have sequence features rather consistent across cell types, and CTCF-MP can be effectively used to predict CTCF loops in a new cell type by using both sequence-based feature and selected functional genomic signals (CTCF ChIP-seq and DNase-seq).

**Figure 4:**
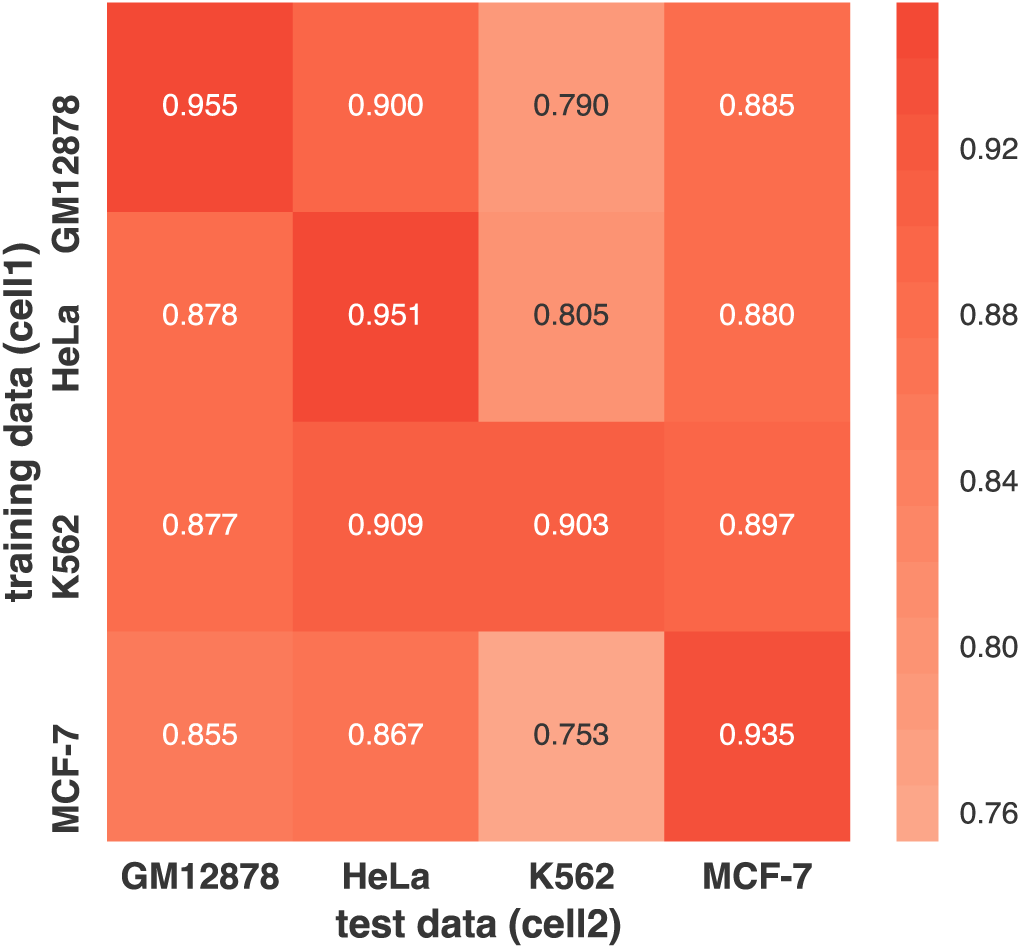
Performance of using CTCF-MP for cross cell type prediction. The number shows the AUROC result for model trained on cell type 1 and tested on cell type 2.

**Figure 5:**
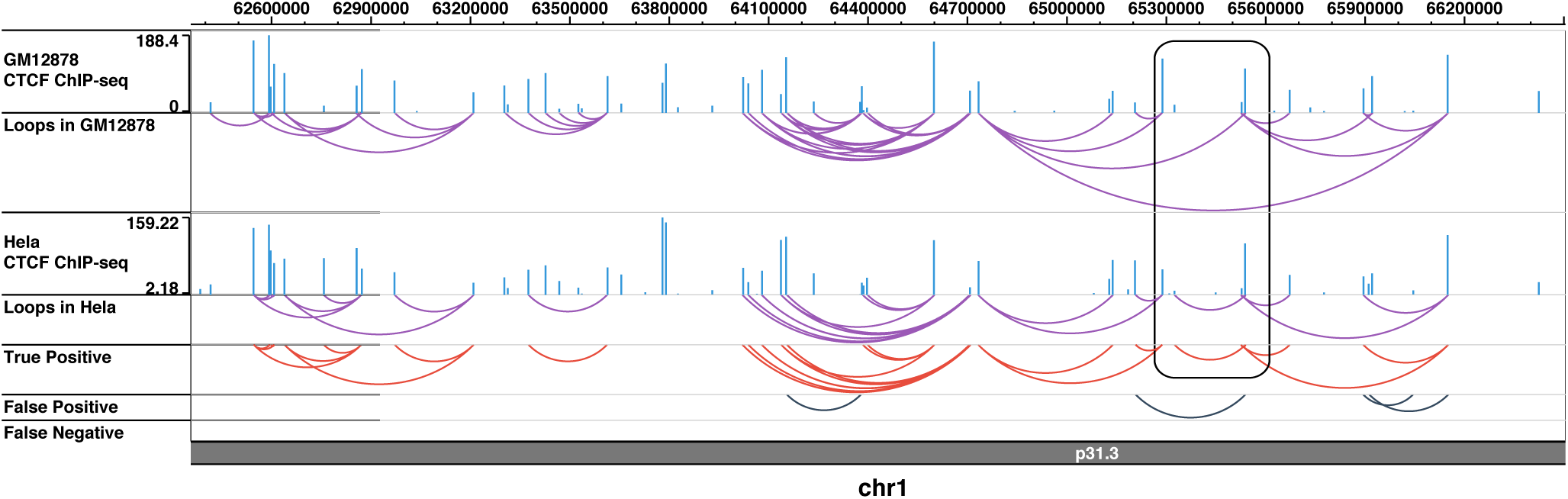
An example of the cross cell type prediction from CTCF-MP. The highlighted box in the figure shows cell type specific loop in HeLa that is correctly predicted by CTCF-MP based on trained model from GM12878 ChIA-PET data. “GM12878/HeLa CTCF ChIP-seq” track shows the ChIP-seq peak value for the CTCF motifs in GM12878 and HeLa, respectively. “Loops in GM12878/HeLa” shows the CTCF loops based on ChIA-PET in GM12878 and HeLa, respectively. “True Positive/False Positive/False Negative” are the predictions in HeLa made by CTCF-MP where the classifier is trained in GM12878.

Furthermore, we asked if the performance can be further improved when we use training data from more than one cell type. We tested this idea by using data from HeLa, K562, and MCF7 as training data to build the classifier and GM12878 as testing cell type. Similarly, we required that there are no shared CTCF pairs in training and testing data in order to only consider cell type specific CTCF loops that appear only in GM12878. CTCF-MP reaches 90.8% AUROC, which is better than the setting where data from only one cell type is used. We also tested in the whole GM12878 CTCF pair dataset (imbalanced, positive : negative = 22,432 : 215,607), after tuning the threshold to reach the highest F1 score. CTCF-MP reaches 88.7% accuracy overall. Detailed results can be found in Table S1. Taken together, the results here demonstrated the potential of CTCF-MP to predict CTCF loops for a new cell type.

### CTCF-MP extracts important features for convergent CTCF motif pairs that form loops

In CTCF-MP, we used word2vec-encoded vector, extra sequence-based features (branch-of-origin, distance between the motif pair, GC content, CTCF motif occurrence frequency, matching score to motif PWM computed by FIMO (see Methods), and functional genomic signals from CTCF ChIP-seq and DNase-seq to train the classifier using different features individually as well as different combinations of features (see Table 2 for detailed results). Although using all features achieves the best performance (AUROC=95.5%), we found that features extracted from word2vec alone can still do well for both AU-ROC and AUPR. As expected, adding extra features in addition to the word2vec features would gradually improve the performance. In particular, we found that, besides word2vec features, the branch-of-origin score itself can achieve good predictive power (AUROC=76.6%) to distinguish loop-forming convergent CTCF motif pairs. When branch-of-origin is combined with CTCF ChIP-Seq signals, we can reach even better performance than using each feature individually, suggesting that branch-of-origin provides more information than CTCF occupancy reflected by the ChIP-seq signal to further distinguish loop-forming CTCF motif pairs. We found that the sequence-based features alone (word2vec + all extra sequence features) can predict CTCF loops based on convergent CTCF motif pairs without using any functional genomic signals with high accuracy. In addition, to further demonstrate word2vec’s ability for encoding DNA sequences, we added comparison between word2vec encoded vectors and traditional *k*-mer frequency (*k*=6 to be consistent with what we used in word2vec). We found that word2vec can reach much better performance than the *k*-mer method. Taken together, these results suggest that the sequence-based features from word2vec and other sequence level features such as branch-of-origin are informative and complementary to CTCF ChIP-seq and DNase-seq to predict CTCF chromatin loops.

**Table 2:** The impact of different features and combinations with word2vec features in predicting loop-forming convergent CTCF motif pairs in GM12878. All extra sequence features include branch-of-origin, distance between motif pairs, GC content, motif occurrence between paired motifs, and matching score to motif PWM.

|  | Accuracy | Precision | Recall | F1 | AUROC | AUPR |
| --- | --- | --- | --- | --- | --- | --- |
| $k$-mer only | 63.0% | 0.658 | 0.540 | 0.593 | 0.680 | 0.647 |
| word2vec only | 70.5% | 0.656 | 0.864 | 0.746 | 0.796 | 0.776 |
| word2vec + all extra seq. features | 80.4% | 0.769 | 0.869 | 0.816 | 0.893 | 0.889 |
| word2vec + all extra seq. features<br>+ ChIP-seq & DNase-seq | 88.8% | 0.869 | 0.915 | 0.891 | 0.955 | 0.951 |
| CTCF ChIP-seq &<br>branch-of-origin | 79.4% | 0.806 | 0.773 | 0.790 | 0.880 | 0.850 |
| CTCF ChIP-seq Only | 77.4% | 0.776 | 0.771 | 0.773 | 0.849 | 0.802 |
| DNase-seq Only | 72.6% | 0.757 | 0.667 | 0.709 | 0.791 | 0.739 |
| ChIP-seq & DNase-seq | 78.6% | 0.787 | 0.786 | 0.786 | 0.863 | 0.826 |
| all extra seq. features only | 77.2% | 0.772 | 0.772 | 0.772 | 0.862 | 0.862 |
| distance only | 54.0% | 0.727 | 0.126 | 0.214 | 0.565 | 0.587 |
| branch-of-origin only | 67.3% | 0.800 | 0.462 | 0.586 | 0.766 | 0.755 |

We further evaluated the importance of each dimension of word2vec features in the classifier. In each round of the 10-fold cross-validation, after training the model with word2vec features only, CTCF-MP estimated feature importance, the information gain of the feature when it is used in trees, and used the average of it as the evaluation criteria. In Fig. 6A, we show the feature importance across different cell types. Although the actual meaning of each dimension of word2vec features is difficult to interpret due to the nature of word embedding, it does show that the most predictive features are generally consistent across different cell types. We visualized the distribution of the samples in the vector space that all our features established. We started by using a deep autoencoder [19] to compress the 200-dimensional space (100 dimensions from each side of the pair) from word2vec together with the other sequence features we used into a lower dimensional space (32-dimensional). Then we applied Stochastic Neighbor Embedding (t-SNE) [20] to map the compressed data into a 2-D plane. In Fig. 6B, x-axis and y-axis form the 2-D plane that t-SNE compressed to, where each point represents a sample with its color representing the label (either forming loop or not forming loop). We found that positive samples and negative samples are clustered mostly together based on the features from word2vec and other sequence-based features, suggesting that our CTCF-MP method constructs an efficient vector space for our classification purpose.

**Figure 6:**
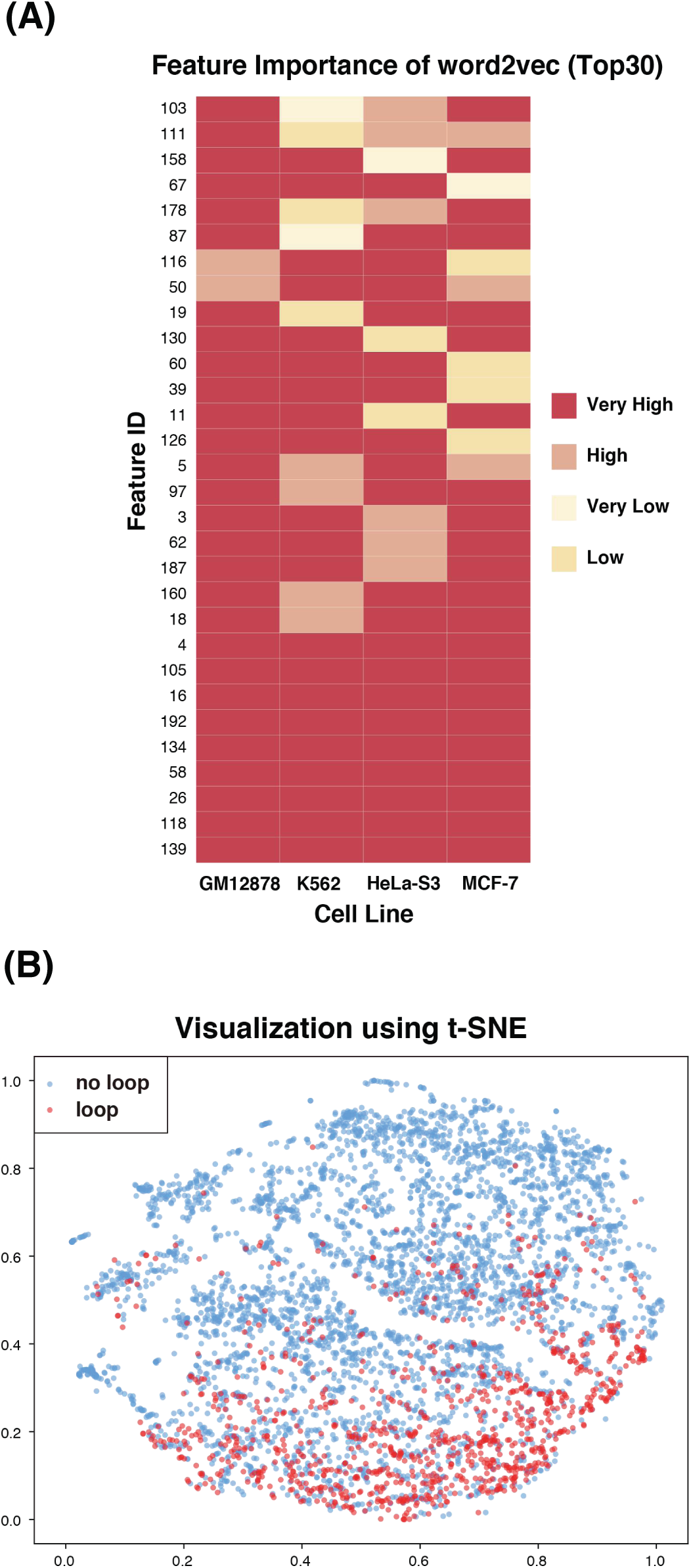
(A) Feature importance of word2vec features across different cell types. (B) Visualization using t-SNE based on word2vec features and other sequence-based features we used in CTCF-MP.

## Methods

### Data collection

We downloaded the CTCF ChIA-PET data on K562 and MCF7 from the ENCODE project website and CTCF ChIA-PET data from GM12878 and HeLa from GEO (accession: GSE72816). We also downloaded CTCF ChIP-Seq peaks and DNase-seq peaks from ENCODE. Mammalian phyloP scores were downloaded from the UCSC Genome Browser. The known human CTCF motif PWM was obtained from JASPAR (ID: MA0139.1) [21]. We used FIMO [22] to scan all CTCF motifs in the human genome. We used the default parameters and a *p*-value 1e-5 as threshold. We overlapped these motifs with long-range interaction peak regions (called from ChIA-PET) to define loop motifs and non-loop motifs.

### The CTCF-MP algorithm

#### Training and testing datasets in CTCF-MP

We defined positive and negative samples for the machine learning module in CTCF-MP as follows. We considered motif pairs (that are bound by CTCF) with convergent orientation only (for both positive and negative cases). For unique CTCF motifs within the ChIA-PET detected loop regions, the motif pairs were considered as positive samples. If there were more than one motif in either side of the paired ChIA-PET loop region, those motifs were not included in the classification (i.e., they were considered neither as positive samples nor as negative ones). In other words, we defined positive samples as the following: in a ChIA-PET defined loop, on either side of the paired regions there is a unique CTCF motif bound by CTCF. We then estimated the distance distribution of positive samples by calculating a range that can cover 95% of the positive samples, which was then used as the distance for generating negative samples. Negative samples were those motif pairs within the distance range but were not in the positive samples, i.e., they did not form loops. We then sampled negative samples with similar distance distribution as positive ones. The main reason of this approach was to minimize the contribution of distance between a pair so that we can focus on understanding other features.

We hypothesized that whether two CTCF motifs could form a chromatin loop depends on both the features they have individually and the features they share. We grouped the negative samples into four categories depending on whether those two CTCF motifs are loop motifs or not. We trained our model with a softmax loss function for multi-classification. But when we evaluated the algorithm, we combined the four negative labels into one and used binary classification metric to evaluate the performance. For all the evaluations, we used 10-fold cross-validation to train and test our method.

#### The word2vec model in CTCF-MP

From the recent development of learning word embedding approaches in the field of natural language processing, word2vec, which uses distributed representation of words in a continuous vector space, has been proven to be an effective method to reduce the high dimensionality of word representations in contexts [13]. Word2vec is a two-layer neural network that learns embedding vectors for words in the text corpus. The main idea is that we can encode words within a text corpus by establishing potential interactions between the word and its contexts to discover important patterns in natural language. In such a model, words are embedded in a continuous vector space where “semantically similar” words have closer vectors. The basic idea for training such a model is that words that appear in the same contexts share semantic meaning. Thus, words and their contexts from the corpus are used as positive samples to train a model through multiple ways. Here we utilized word2vec to train a distributed representation and encoding for DNA sequences. We recently used a similar approach to predict enhancer-promoter interactions [23]. We considered subsequences of fixed length *k* as DNA “words” (also referred to as *k*-mers). The collection of all possible *k*-mers was defined as the vocabulary (size of vocabulary = 4*^k^*). We then used a *k* sized sliding window to scan sequence with the CTCF motif as well as its flanking region with step size 1 to build a DNA “sentence”.

After we built DNA sentences based on the CTCF motifs and their flanking regions, we used them as training data for word2vec to build a Continuous-Bag-of-Words (CBOW) model [24] for establishing a representation vector space for DNA sequences. A CBOW model aims to predict a word from its neighbors, and the parameters of the model can be presented as a matrix of *V* × *N*, where *V* is the size of vocabulary and *N* is the dimensionality of the embedded feature space. After we trained the model, each row of the learned parameters can be regarded as the embedding vector for a specific *k*-mer word. The probabilistic model for this problem is trained by maximizing the probability of target word *w*_*t*_ given the context *c*, i.e.,

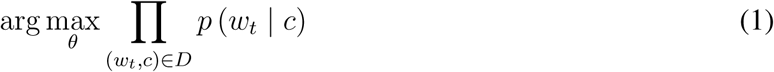

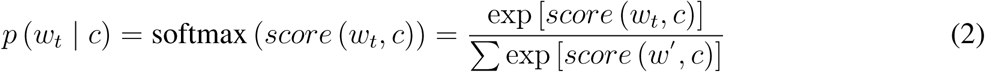

where *D* is the set of all pairs of word and context in the sequences, *score* (*w*_*t*_, *c*) computes the compatibility of words (e.g., using a dot product) and *w*^′^ represents all possible words in the vocabulary. However, such a language probabilistic model is computationally very inefficient. Word2vec uses a technique called negative sampling [25], which trains a binary classification model to discriminate the real target word *w*_*t*_ from “noise” words 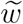, given the same context *c*. Noise words are sampled from noise distribution estimated from the text corpus (in our case, sequences). In other words, positive samples are those pairs of word and context that have appeared in the sequences while negative samples are those that have not. The model is trained by maximizing an objective function that achieves higher score when the model assigns high probabilities to the real words and low probabilities to noise words, i.e.,

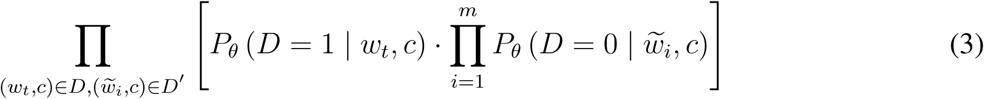

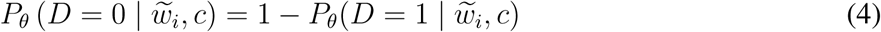

where *D* is the set of all pairs of word and context in the positive samples, *D*′ is the set o negative samples, and *P*_*θ*_ (*D* = 1 | *w*_*t*_, *c*) is the probability that the word and context pair (*w*_*t*_, *c*) is observed in positive samples for the learned parameter vector *θ*. The objective function scales only with the number of *m* noise words instead of all words in the vocabulary.

After training the word2vec model, we have the embedding vectors for each DNA word. To put it in another way, we can have a hash table with its keys as DNA words and its values as vectors. We then encoded DNA sequences by having the embedding vectors for every DNA word in the DNA sentence and taking the average of the vectors as the vector for the DNA sequence.

In CTCF-MP, we set *k* = 6 as the word length, *N* = 100 to be the dimensionality of the embedded features, and +/- 250 bp of the motif as the flanking region, by balancing the amount of sequence patterns we would like to model and computational cost.

#### Additional features in CTCF-MP

After encoding DNA sequences into vectors, we selected other features based on some prior knowledge and our own observations (e.g., branch-of-origin of CTCF motifs). To capture the sequence-based information on whether a CTCF motif has the ability to form loops, we included the following features: branch-of-origin of CTCF motifs, CTCF motif matching score to the motif PWM, GC content, distance between motif pairs, motif occurrence in the genomic region between CTCF motif pair. In addition, we also included signals from CTCF ChIP-seq and DNase-seq of the regions under consideration.

#### Boosted trees classifier in CTCF-MP

In the classification step, we considered all the features from word2vec modeling step together with the additional features as input for a boosted tree classifier. Like other ensemble learning methods, boosting aims to combine a set of weak learners to be a stronger classifier [26]. The core idea of boosting is to iteratively train models that add more weight to the misclassified samples and thus ultimately achieve a better classifier. A decision tree is typically used as the weak learner in boosting algorithms and has both great performance and high efficiency.

In CTCF-MP, we used the gradient boosting algorithm [27]. For this multi-classification setting, the algorithm tries to minimize the softmax loss function. In each iteration stage of gradient boosting, it improves the existing model by adding an extra estimator to it. The process repeats until it reaches the maximum iteration rounds or convergence. For this 5-class problem (as discussed earlier, one positive type and four negative types) with training set {(*x*_1_,*y*_1_), …, (*x*_*n*_, *y*_*n*_)},*y*_*i*_ ∈ {0,1, 2, 3, 4}, the loss function of iteration stage *m* is as follows:

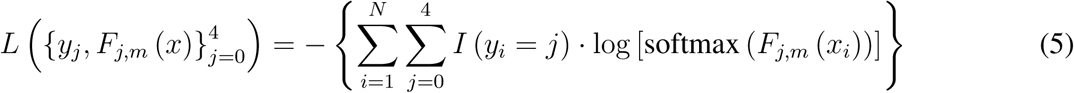

where

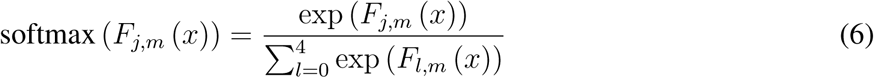

where *F*_*j,m*_ represents the learned model for class *j* on stage *m*. *I* (*y*_*i*_ = *j*) is the characteristic function that equals 1 when *y_i_ = j*. In step *m*, the algorithm would fit five decision trees *h*_*j,m*_ (*x*), *j* = 0, …, 4 to predict residuals for each class on the probability scale. If each tree has *K* nodes, with corresponding regions {*R*_*kjm*_},*k* = 0, …, *K –* 1, j = 0, …, 4, the model updates as follows:

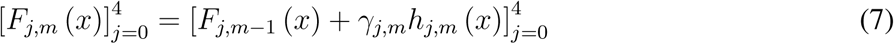

where

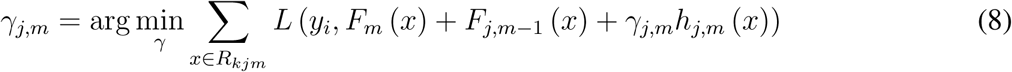

and

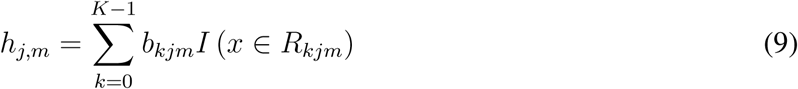

where *b*_*kjm*_ is the value predicted in region *R*_*kjm*_. In CTCF-MP, we used XGBoost [28], which is an excellent boosting implementation. XGBoost can train the model by multi-thread operation and has rather high performance and robustness to over-fitting.

### Method to calculate Branch-of-Origin

For each CTCF binding motif occurrence, we obtained +/-100bp orthologous sequence centered on the human CTCF binding site across mammalian species using the Multiz alignment available on UCSC genome browser. Next, motif occurrence in 200bp multiple sequence alignment block across different species were counted. We then applied the birth-death model initially described in [14] to predict the branch-of-origin of each CTCF motif occurrence in human. See [14] for the details of the method that models cis-regulatory element evolution.

## Discussion

In this work, we developed effective computational methods to address several important questions related to CTCF-mediated chromatin loops. One of our main motivations is to evaluate the contributions of sequence-based features already encoded in the genome that may provide instructions to determine CTCF chromatin loops. Our results allow us to answer the three questions we proposed at the beginning:

a. We found that motif conservation measured by “branch-of-origin” that accounts for motif turn-over in evolution is an informative feature to distinguish loop motifs from non-loop motifs.
b. For an individual cell type, we can train a CTCF-MP classifier (based on word2vec and boosted trees) to accurately predict loops formed by convergent CTCF motifs bound by CTCF using both sequence features as well as CTCF ChIP-seq and DNase-seq. In particular, we found that sequence-based feature alone have strong capability to predict if a pair of convergent CTCF motifs would form a loop.
c. We can train a CTCF-MP classifier based on data from existing cell type(s) to effectively predict whether a pair of convergent CTCF motifs would form a loop (including cell type specific loops) in a new cell type.

Our work offers important new insights in the sequence-based features underlying loop formation between a pair of CTCF motifs. In the recent work from [12], the authors found that epigenetic marks together with CTCF motif occurrences can be used to predict chromatin loops between a pair of convergent motifs. However, there are several main differences in our work: (1) In this paper, we focus mainly on the contributions of sequence-level features in forming loops. Our work demonstrated CTCF-MP’s potential to predict CTCF loops for cell types without many functional genomic datasets. (2) It is known that the majority of CTCF loops have convergent motif orientation and the distance is one of the most discriminative features in deciding whether CTCF motif pairs would form a loop. [12] did not specifically consider this factor. We carefully prepared the data to reduce the contribution of distance to the model, such that we can discover other more important and novel features.

There are a number of areas that our methods and approaches can be further improved to reveal a more complete picture of CTCF-mediated chromatin loops. For example, at the moment, we focus on convergent CTCF motif pairs as those are the ones that have been consistently observed in both Hi-C data and ChIA-PET data. However, in [8], the authors reported that in addition to convergent pairs there are also about 33% of motif pairs among detected CTCF loops that are “in tandem”. We have made initial evaluation on CTCF-MP’s performance to predict loops formed by tandem CTCF motifs (see Supplementary Results and Tables S2-S3). It would be useful to further explore the differences in sequence features of the pairs in tandem and compare with convergent ones and understand their functional importance. In addition, one limitation in the methodology of our CTCF-MP algorithm now is that the features from word2vec are hard to interpret due to the difficulty in clearly explaining the embedded space (in fact, the same challenge also exists in the field of natural language processing even though word2vec has been successfully applied in NLP) [25]. Usually, visualization algorithms such as t-SNE can be used to provide an idea of the embedded space. Nevertheless, our evaluation demonstrated that our word2vec features alone can already predict loop-forming convergent motif pairs with quite good performance. It is also encouraging that with the additional sequence-based features (such as branch-of-origin) that are not captured by word2vec model, CTCF-MP achieves high performance in prediction without functional genomic signals from ChIP-seq and DNase-seq. Overall, we believe our methods and results made an important step further in our understanding of the principles of CTCF-mediated chromatin loops. CTCF-MP could be particularly useful when we prioritize and interpret mutations in human disease genomes (e.g., for a better understanding of somatic non-coding mutations in tumor genomes). The insights from our work also have the potential to help decode information encoded in our genome sequences that determine complex chromatin architectures.

## Acknowledgments

This work was supported in part by National Institutes of Health grants R01HG007352 and U54DK107965 (J.M.), and National Science Foundation grants 1054309 and 1262575 (J.M.). R.Z. was previously supported by Tsinghua University’s Top Open program.

